# A chromosome-scale genome assembly for the *Fusarium oxysporum* strain Fo5176 to establish a model *Arabidopsis*-fungal pathosystem

**DOI:** 10.1101/2020.05.07.082867

**Authors:** Like Fokkens, Li Guo, Susanne Dora, Bo Wang, Kai Ye, Clara Sánchez-Rodríguez, Daniel Croll

**Affiliations:** Molecular Plant Pathology, University of Amsterdam, The Netherlands; MOE Key Laboratory for Intelligent Networks & Network Security, Faculty of Electronic and Information Engineering, Xi’an Jiaotong University, Xi’an, China; Department of Biology, ETH Zurich, Zurich, Switzerland; Laboratory of Evolutionary Genetics, Institute of Biology, University of Neuchâtel, Neuchâtel, Switzerland

**Keywords:** Fusarium oxysporum, Arabidopsis thaliana, genome assembly, transposable elements, infection assays, comparative genomics

## Abstract

Plant pathogens cause widespread yield losses in agriculture. Understanding the drivers of plant-pathogen interactions requires decoding the molecular dialog leading to either resistance or disease. However, progress in deciphering pathogenicity genes has been severely hampered by suitable model systems and incomplete fungal genome assemblies. Here, we report a significant improvement of the assembly and annotation of the genome of the *Fusarium oxysporum* (*Fo*) strain Fo5176. *Fo* comprises a large number of serious plant pathogens on dozens of plant species with largely unresolved pathogenicity factors. The strain Fo5176 infects *Arabidopsis thaliana* and, hence, constitutes a highly promising model system. We use high-coverage Pacific Biosciences Sequel long-read and Hi-C sequencing data to assemble the genome into 19 chromosomes and a total genome size of 67.98 Mb. The genome has a N50 of 4 Mb and a 99.1% complete BUSCO score. Phylogenomic analyses based on single-copy orthologs clearly place the Fo5176 strain in the *Fo* f sp. *conglutinans* clade as expected. We generated RNAseq data from culture medium and plant infections to train gene predictions and identified ∼18,000 genes including ten effector genes known from other *Fo* clades. We show that Fo5176 is able to infect cabbage and Brussel sprouts of the *Brassica oleracea*, expanding the usefulness of the Fo5176 model pathosystem. Finally, we performed large-scale comparative genomics analyses comparing the Fo5176 to 103 additional *Fo* genomes to define core and accessory genomic regions. In conjunction with the molecular tool sets available for *A. thaliana*, the Fo5176 genome and annotation provides a crucial step towards the establishment of a highly promising pathosystem.

## Introduction

*Fusarium oxysporum* (*Fo*) is an important threat to food production worldwide infecting more than 100 different crops (Edel-Hermann & Lecomte 2019). Soil-borne vascular pathogens colonizing plants through the roots make chemical, cultural and biological controls generally ineffective. To tackle pathogen threats durably, suitable model pathosystems and deep molecular knowledge of the infection process are essential. Individual *Fo* strains have a narrow host specificity, on which strains can be grouped into distinct *formae speciales*. The *Arabidopsis thaliana – Fo* pathosystem offers a wealth of tools and resources including genotyped plant accessions mutant. This makes the system an ideal model to generate fundamental insights into how *Fo* establishes infections and how plant resistance can be improved. To develop the pathosystem into an efficient model, high-quality genome assemblies and accurate gene annotations of *Fo* strains that infect Arabidopsis have been lacking. The Fo5176 strain, originally isolated in Australia from *Brassica oleracea* plants (Thatcher et al. 2012; Chen et al. 2014), is virulent on multiple *Arabidopsis* accessions (Thatcher et al. 2009). The currently available Fo5176 genome assembly published in 2011 lacks significantly in contiguity (Thatcher et al. 2012). Yet high-quality genome assemblies are crucial due to the abundance of repetitive sequences surrounding key pathogenicity-related genes. Comparative genomic studies have revealed that the genome of *Fo* consists of 10-11 core chromosomes and a variable number of accessory chromosomes (Williams et al. 2016; Armitage et al. 2018; van Dam et al. 2017; Ma et al. 2010)). In tomato and cucurbit-infecting strains pathogenicity genes are clustered on a subset of the accessory chromosomes, known as pathogenicity chromosomes. Horizontal transfer of these chromosomes can contribute to the emergence of new pathogenic strains (van Dam et al. 2017; Li et al. 2020; Ma et al. 2010).

Here, we present the assembly and annotation of the Fo5176 genome to complete the *Arabidopsis* - *Fo* pathogenicity model. Based on PacBio SMRT (single-molecule real-time) and Hi-C (high□throughput chromosome conformation capture) sequencing technologies, we achieved a robust chromosome-level assembly of the genome. We used high-quality RNAseq data to train the gene annotation and predicted about 18,000 genes. In a comparative genomics analysis including a large number of additional *Fo* genome sequences, we identify regions putatively determining host preference within the species. Finally, we assayed the infection potential of Fo5176 beyond *A. thaliana* on a number of different crop species.

## Materials and methods

### Fungal cultivation and DNA extraction

The Fo5176 strain was originally isolated in Australia as described in (Thatcher et al. 2012). To obtain spores for fungal infection, 400 µl of 10^7^ spores/ml were dark grown for 5 days in 100 ml potato dextrose broth (Laboratorios CONDA) at 27° C at 100 rpm. Spores were filtered through miracloth, centrifuged for 5 minutes at 3750 g and washed twice with sterile water. The non-filtered mycelia were used for DNA extraction. The spore concentration was evaluated on a Thoma counting chamber. For each infection replicate fresh spores were obtained from the same original stock of Fo5176 spores. Genomic DNA was extracted from cultures described above starting with slightly dried mycelia split into 200 µg aliquots and flash-frozen in liquid nitrogen. Then, we followed the method from (Allen et al. 2006) using mortar and pestle to grind the material. We washed the pellet in step 18-19 three times before drying the extracted genomic DNA in a Speedvac for 15 minutes. The DNA was resuspended in 25 µl dH_2_O overnight at 4° C. The integrity of the genomic DNA was inspected by gel electrophoresis on a 1% agarose gel.

Procedures for soil infection assays on *Brassica oleracea* are available in Supplementary Methods.

### Genome assembly and Hi-C analyses

Sequencing was performed at the Functional Genomic Center of Zurich using the PacBio Sequel platform. Eight μg of DNA was mechanically sheared to an average size of 20-30 kb using a g-Tube device (Covaris). The SMRTbell was produced using the SMRTbell Express Template Prep Kit. After size selection to enrich fragments >17 kb, we sequenced the library on 1 Sequel™ SMRT® Cell taking 1 movie of 10 hours per cell. For genome assembly, we used HGAP 4 included in the SMRTlink software version 6.0.0.47841 (Chin et al. 2013). We used three seed length cutoffs for read correction (30, 35 kb and auto-determined at 38.7 kb). The estimated genome size was set to 60 Mb. The assembled contigs were polished using Quiver as implemented in the SMRTlink HGAP 4 pipeline. The three assemblies produced by the different seed lengths were inspected for differences in contiguity using blastn (Altschul et al. 1990) with transcript sequences of the *F. oxysporum* reference genome FO2 (Ma et al. 2010).

To verify and produce a chromosome-level genome assembly, Hi-C sequencing data was generated. Hi-C library construction of *F. oxysporum* Fo5176 was prepared from cross-linked chromatins of fungal cells using a standard Hi-C protocol (Belton and Dekker 2015). The Hi-C library was checked for valid interaction read pair ratios in a test Illumina sequencing run. The quality-controlled library was sequenced by Illumina NovaSeq 6000 to yield 5.10 Gb (∼75x coverage) paired-end reads. The Hi-C sequencing data was used to anchor all contigs using Juicer v. 1.5 (Durand et al. 2016a), followed by using a 3D-DNA correction pipeline (Dudchenko et al. 2017) and manually refined with Juicebox v. 1.11.08 (Durand et al. 2016b).

### RNA extraction and sequencing

Fungal RNA was extracted from *in vitro* and *in planta* Fo5176. To obtain *in vitro* produced spores, 10^7^ spores per ml were germinated overnight in ½ Murashige and Skoog Basal Medium (MS; Sigma-Aldrich), 1% sucrose and kept at 100 rpm. Spores were harvested, washed with sterile water via two centrifugation steps at 4000 g for 15’ at 10° C. The pellet was frozen in liquid nitrogen and subsequently freeze-dried before RNA extraction. *In planta* Fo5176 RNA was obtained from hydroponic-grown *A. thaliana* roots two days after fungal inoculation. Frozen infected roots and germinated spores were ground with mortar and pestle in liquid nitrogen. In total, 50 -100 µg of ground material was used for RNA extraction using the RNeasy plant mini Kit (Qiagen). Total mRNA-Seq libraries were prepared using the SENSE mRNA-Seq library Prep Kit V2 (Lexogen) according to the instruction manual with few modifications. Sequencing was performed in an Illumina HiSeq2500 sequencer (single-end 125 bp) with a read depth of around 120 mio reads per sample. A more detailed protocol of the RNA extraction and sequencing steps is available in Supplementary Methods.

RNA-seq reads were mapped to the assembled genome using HISAT2 version 2.1.0 (Ma et al. 2010; Kim et al. 2019). Mapped reads were merged using samtools (Li et al. 2009). We used braker version 2.1.5 (Hoff et al. 2019) to predict gene models with the “--fungus” option. Genes overlapping contig ends were removed. We used exonerate in the protein2genome mode (Slater & Birney 2005) to localize confirmed effector genes from other *F. oxysporum* genomes (*Six1, Six4, Six8, Six9a, Foa1-4, Foa6 and FoaXY1;* Tintor et al. 2020). Matching regions were added as additional gene models to the annotation. We used InterProScan version 5.31.70 (Slater & Birney 2005; Jones et al. 2014) to functionally annotate gene models. In addition, we analyzed predicted protein sequences for evidence of secretion signals and transmembrane domains using SignalP version 4.1(Petersen et al. 2011) and TMHMM v. 2.0 (Krogh et al. 2001) (Supplementary Table S1). We analyzed the repeat content of the genome using RepeatModeler v. 1.0.8 (http://www.repeatmasker.org/RepeatModeler; Hubley R, Smit AFA). This pipeline includes RECON and RepeatScout to detect repeat families *de novo* and identify consensus sequences. Consensus sequences were manually inspected for consistency in the assignment. Both newly identified repeat families and the RepBase content were then used to annotate the genome using RepeatMasker version 4.0.7 (http://www.repeatmasker.org; Smit AFA, Hubley R & Green P).

### Phylogenomics analyses

We used BLAST (megablast -evalue 0.0000001 –outfmt 6) to search for homologs of all predicted Fo5176 gene sequences (including introns) against a database of 103 *Fo* genomes and the genome of *F. fujikuroi* as an outgroup (Table S3). Individually added effector genes were not included. Genomes were downloaded from Genbank on January 25^th^ 2019 (Supplementary Table S3). We selected hits with more than 80% overlap with the query sequence, selected queries with one selected hit per genome in the dataset and wrote these queries with their selected hits to a fasta file using a custom Python script. We used MUSCLE (Edgar 2004) with default settings to align the sequences in the 6833 fasta files thus obtained. We used TrimAl (Capella-Gutiérrez et al. 2009) to trim the alignments, removing positions with gaps in 10% or more of the sequences, unless this would leave less than 70% of original alignment, in which case we kept the 70% positions with the least amount of gaps (-gt 0.9 - cons 70), where we averaged the gap score over a window starting 3 positions before and ending 3 positions after each column (-w 3). We concatenated the trimmed alignments and created a partition file stating the positions of individual genes in the concatenated alignment using a custom Python script. We used IQTree (version 1.6.12) to find the best substitution model for each partition (-m MFP) and infer a consensus tree using maximum likelihood and ultrafast bootstrapping (*n* = 1000) sampling sites per gene (-B 1000 --sampling GENESITE) (Nguyen et al. 2015; Kalyaanamoorthy et al. 2017).

### Whole-genome alignments and presence-absence analyses

We used nucmer (--max-match) from the MUMmer package version 3.23 (Kurtz et al. 2004) to align the 103 *Fo* genome assemblies (see above) to our assembly of Fo5176. To classify genomic regions into ‘core’ or ‘accessory’, we filtered for each of the 103 *Fo* assemblies the MUMmer output to obtain global alignments (‘deltafilter –g’), wrote alignments that spanned at least 1 kb in Fo5176 to a bed formatted file and used ‘bedtools genomecov -bga -split –i’ (Quinlan & Hall 2010) to calculate the ‘coverage’ (number of genomes that align) per base pair. All base pairs that had ‘coverage’ >= 93 (*i*.*e*. aligned with at least 90% of the genomes in the dataset) were assigned as core regions, the remainder was considered being accessory regions.

### Data availability statement

This Whole Genome Shotgun project has been deposited at DDBJ/ENA/GenBank under the accession JACDXP000000000. The version described in this paper is version JACDXP010000000. Hi-C sequencing data of Fo5176 are available in the NCBI Sequence Reads Archive (SRA) under the accession number SRR11742970.

## Results and Discussion

### Chromosome-scale genome assembly of Fo5176

The *Fo* strain Fo5176 has well-established virulence on *A. thaliana* but lacks a highly contiguous genome assembly (Thatcher et al. 2012). Based on high-coverage PacBio Sequel reads, we assembled the genome into 19 chromosome-scale contigs. We used Hi-C data to correct one inversion and two fusions. The number of chromosomes was determined by centromeric interaction regions detected in Hi-C contact map figures (Marbouty et al. 2014) resulting in a final assembly of 19 chromosomes with a N50 of 4.09 Mb and a genome size of 67.98 Mb (Figure 1A). Several contigs are enriched for telomeric repeats at their edges. Contigs 1, 8, 9 and 18 are enriched in telomeric repeats on both ends indicating that they represent a telomere-to-telomere assembly of chromosomes (Figure 1C). Compared to most known *Fo* genomes, the Fo5176 genome is very large and 13.2 Mb larger than the current publicly available assembly of the same strain Fo5176 (GCA_000222805.1). The Fo5176 genome assembly is highly complete with 99.1% complete BUSCO genes using a Sordariomyceta database. A total of 97.1% of the BUSCO genes were single-copy and complete, 1.1% were duplicated, 0.8% were fragmented and 0.4% were missing. The genome has an overall GC-content of 48.2% and a transposable element content of 22.2%. All contigs contain a single, extremely AT-rich region that likely corresponds to the centromere (Figure 1D). The most abundant characterized TEs were DNA transposons (8.79%), long terminal repeats (LTR; 1.75%) and long interspersed nuclear elements (LINE; 1.42%; Supplementary Table S2). Unclassified elements accounted for 10.22% of the genome. The content in TEs is highly variable among chromosomes with markedly higher TE densities on some chromosomes (Figure 1F).

**Figure 1:**
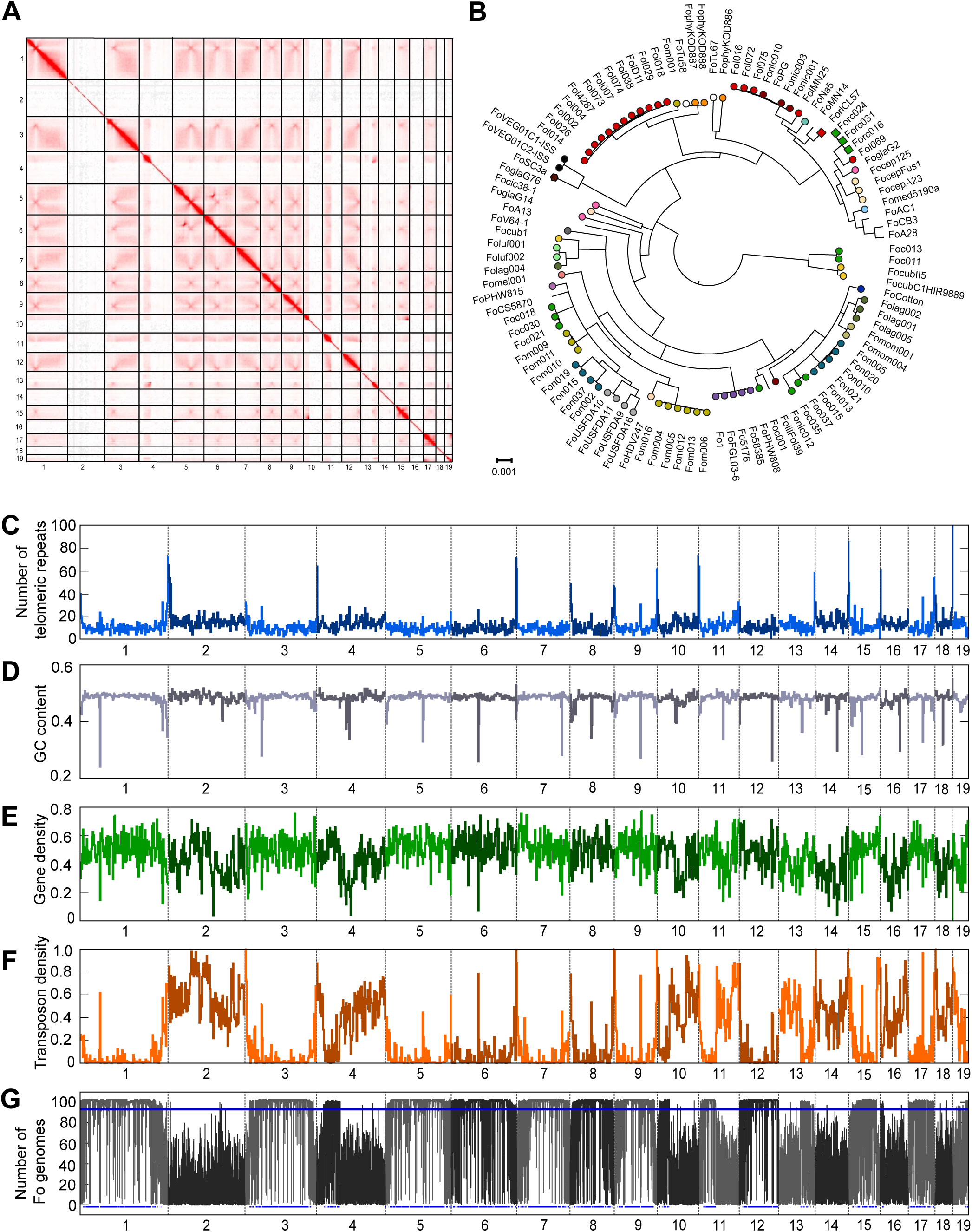
Complete genome assembly of the *Fusarium oxysporum* (*Fo*) strain Fo5176. A) Hi-C contact map of the corrected genome assembly (chromosome 1-19). B) Phylogenomic tree of 103 *Fo* strains (Table S3). Colors indicate formae speciales. Genome scans for C) the number of telomeric repeats, D) GC content, E) gene density, F) repeat density and G) sequence conservation within the species as indicated by the number of *Fo* genomes sharing a chromosomal segment (103 *Fo* strains in total). The bottom blue lines indicate Fo5176 regions that span more than 10 kb and are classified as ‘core’ regions among *Fo* genomes.

Based on RNA-assisted gene annotation taking into account evidence for transcription both in culture medium (*in vitro*) and during infection, we identified 17’902 genes in the Fo5176 genome. We manually added previously known *F. oxysporum* effector gene homologs (*Six1, Six4, Six8, Six9a, Foa1-Foa4, Foa6* and *FoaXY1;* Tintor et al. 2020) to the annotation as these effector genes were either not identified by our gene prediction pipeline or the gene model was no exact match, bringing the total to 17’912 genes. Gene density was largely similar among all chromosomes even on chromosomes lacking homologs with FO2 (Figure 1E). Predictions of proteins with secretion signals, transmembrane domains and conserved domains are provided in Supplementary Table S1. In order to robustly place the Fo5176 genome in the context of the *Fo* diversity, we performed a phylogenomic analysis based on orthologous genes. Previously *Fo* f sp. *conglutinans* was assumed to be a monophyletic group (Enya et al. 2008; Liu et al. 2019). We generated a concatenated alignment of 6833 genes conserved in 103 publicly available *Fo* genomes and the genome of *F. fujikuroi* (Supplementary Table S3). We find that Fo5176 groups with the four other *Fo* f. sp. *conglutinans* strains in a single, well-supported clade unlike other, more densely sampled *formae speciales*, such as *lycopersici* (tomato), *cubense* (banana), or *melonis* (melon) that are polyphyletic (Figure 1B).

### Fo5176 is a pathogen of Brassica oleracea species

A previous study has shown that Fo5176 is capable to infect cabbage plants (Li et al. 2016). To further explore whether Fo5176 may serve as a model pathogen beyond the *A. thaliana* system, we tested whether *Brassica oleracea* species can become infected. *B. oleracea* includes important food crops such as cabbage (*B. oleracea var. capitata*, cv ‘Shikidori’) and Brussels sprouts (*B. oleracea var. gemmifera*, cv ‘Roem van Barendrecht’). We compared the virulence of Fo5176, *Fo* f. sp. *conglutinans* PHW808 (NRRL 54008, hereafter FoPHW808) and the positive control *Fo* f. sp. *conglutinans* Cong: 1-1 (FoCong: 1-1), known to be able to infect *B. oleracea var. capitata, cv ‘Shikidori’* (Kawabe et al. 2011). Both the strain Fo47 (NRRL 54002), used as a biocontrol and not known to be pathogenic on plants (Alabouvette et al. 1986) and *Fo* f. sp. *raphani* PHW815 (NRRL 54005 FoPHW815) infecting related *Brassica* species (*Raphanus sativus* or radish) were used as negative controls. As expected, no disease symptoms were observed in plants inoculated with FoPHW815 or Fo47, while FoCong: 1-1 reduced the weight and generated disease symptoms in both plants (Supplementary Figures S1 and S2). Interestingly, both plant species showed disease symptoms and weight reduction upon infection with both Fo5176 and FoPHW808, indicating that these strains are virulent on both crops (Figure 2A-B; Supplementary Figures S1 and S2) and confirming previous results of Fo5176 virulence in cabbage. Our data shows that the Fo5176 infection model can be expanded to study pathogenicity on *B. oleracea* species.

**Figure 2:**
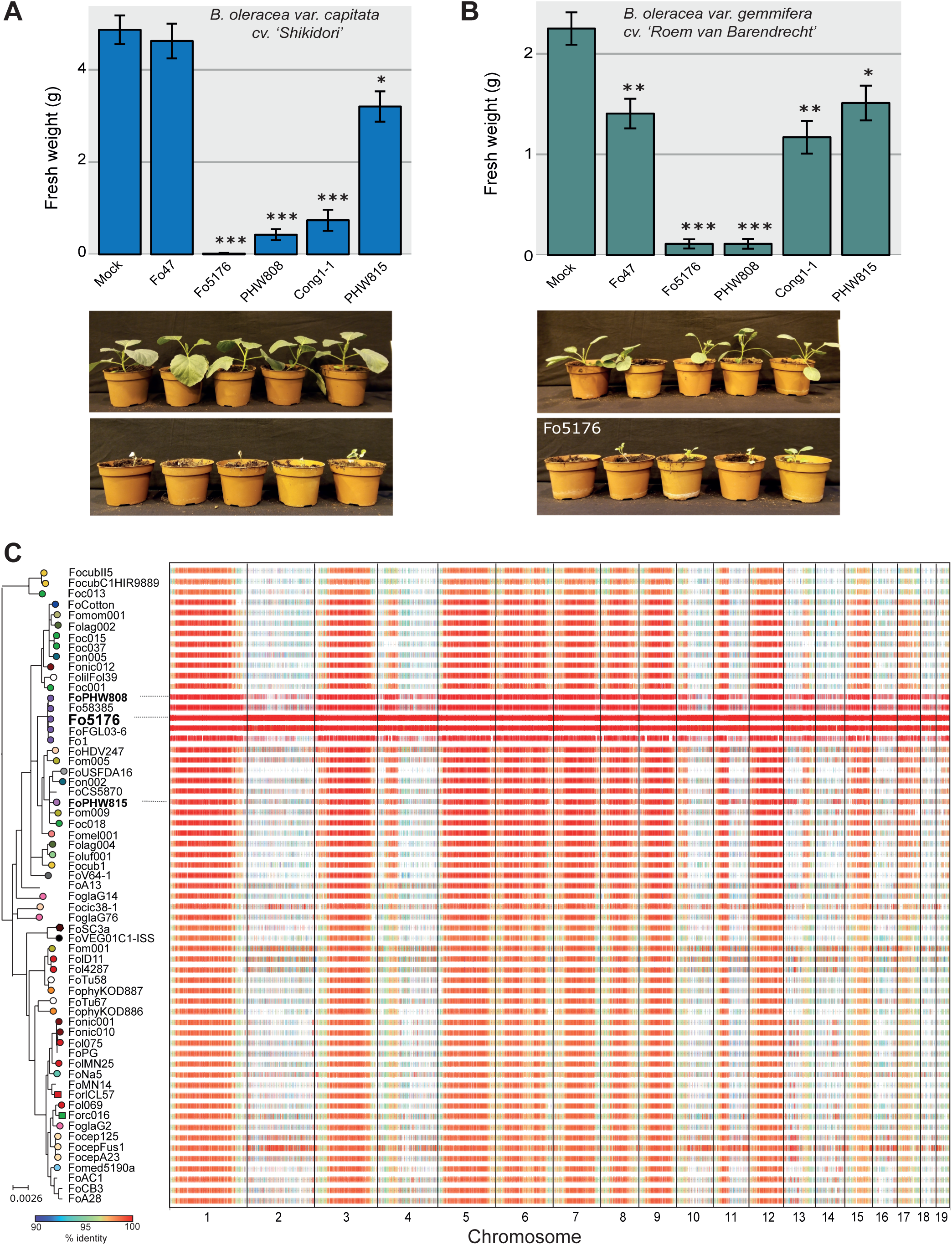
Pathogenicity profiles and sequence conservation among lineages of the broad host range pathogen *Fusarium oxysporum* (*Fo*). A) Infection assay of representative *Fo* strains including Fo5176 on cabbage (*Brassica oleracea var. capitata* cv. ‘Shikidori’). Photographs show Fo5176 and mock-infected plants 13 days after treatment. B) Infection assay with the same *Fo* strains on Brussels sprouts (*B. oleracea var. gemmifera* cv. ‘Roem van Barendrecht’). A more detailed description of both infection experiments is shown in Supplementary Figures S1 and S2. C) Conservation of Fo5176 genomic regions in 103 representative *Fo* genomes. Alignments that span more than 1 kb are more than 90% identical are shown and colored according to percent identity.

### Landscape of the accessory genome across Fo

*Fo* genomes consist of distinct ‘core’ (present in most *Fo* assemblies) and ‘accessory’ (absent in most *Fo* assemblies) regions (Fokkens et al.). We map these regions in the Fo5176 genome, we aligned the new assembly against 103 other *Fo* whole-genome assemblies (Figure 2C; Supplementary Table S3). We classified regions as ‘core’ or ‘accessory’ based on the number of *Fo* genomes the region is aligned to. We find that only 53% of the 67.9 Mb Fo5176 genome is conserved in most other *Fo* genomes and thus considered as ‘core’ regions. These conserved regions are mostly located on chromosomes 1, 3, 5-9, 12, 15 and 17 (Figure 2C). Interestingly, chromosomes 4, 10 and 13 contain large core regions fused to large accessory regions. The remaining contigs consist mainly of lineage-specific accessory regions (Figure 2C). Accessory regions are enriched in TEs and largely depleted in protein-coding genes (Figure 1F-G, 2C). In the tomato-infecting strain Fol4287, partial duplications and triplications of the accessory genome have inflated the genome size (Ma et al. 2010). We did not observe any large-scale duplications in either the core or accessory genome of Fo5176. Hence, the expanded genome of Fo5176 is more likely to be caused by TE proliferation than large-scale duplications.

*Fo* strains infecting tomato or melon express effector genes that are associated with pathogenicity towards a specific host and these effector genes are found on distinct chromosomes (Li et al. 2020; van Dam et al. 2017; Armitage et al. 2018; Ma et al. 2010). Horizontal chromosome transfer can transform a non-pathogen recipient into a pathogenic strain on the host of the donor strain. Horizontal transfer is an important mechanism driving the evolution of pathogenicity in *Fo* (Ma et al. 2010; van Dam et al. 2017; Li et al. 2020). Mobile chromosomes can readily be identified using comparative genomics by testing for matches in the phylogeny of individual chromosomes and hosts. However, such analyses will not allow to distinguish accessory regions conserved among *forma specialis conglutinans* strains because of the shared vertical descent and accessory regions sharing high similarity because of a recent horizontal chromosome transfer event. To select candidate genes involved in host-specific infection processes, gene expression analyses during host infection or screens for protein-protein interactions should be particularly fruitful. The complete genome assembly and high-quality gene annotation presented in this work will lay the foundation for large-scale candidate screens and follow-up studies characterizing the role of individual proteins. The complete genome will provide accurate flanking sequences of targeted genes and ensure that genes embedded in repeat-rich accessory regions are not missed in screens. Finally, the expansive molecular and genetic toolbox available for investigating *A. thaliana* gene functions will enable rapid progress to decipher *Fo* pathogenicity factors.

## Acknowledgements

We thank Prof. T. Arie for providing the FoCong 1-1 strain and seeds for *B. oleracea var. capitata* cv. ‘*Shikidori’*. We acknowledge the technical support of Gloria Sancho and Andrea Patrignani from the Functional Genomic Center Zurich (FGCZ). We also thank Ludek Tikovsky and Harold Lemereis for support in the green house, Tieme Helderman for relabeling inoculum and Sophie Overbeek for assistance in the *Brassica oleracea* infection assay. LF is financially supported by an NWO Veni grant (016.veni.181.090). CSR was supported by the Swiss National Foundation (SNF 310030_184769). LG and KY are supported by the National Natural Science Foundation of China (31701739 and 31671372) and National Key R&D Program of China (2018YFC0910400).

