## Supplementary Information for "A chromosome-scale genome assembly for the *Fusarium oxysporum* strain Fo5176 to establish a model *Arabidopsis*-fungal pathosystem"

### Supplementary Figures

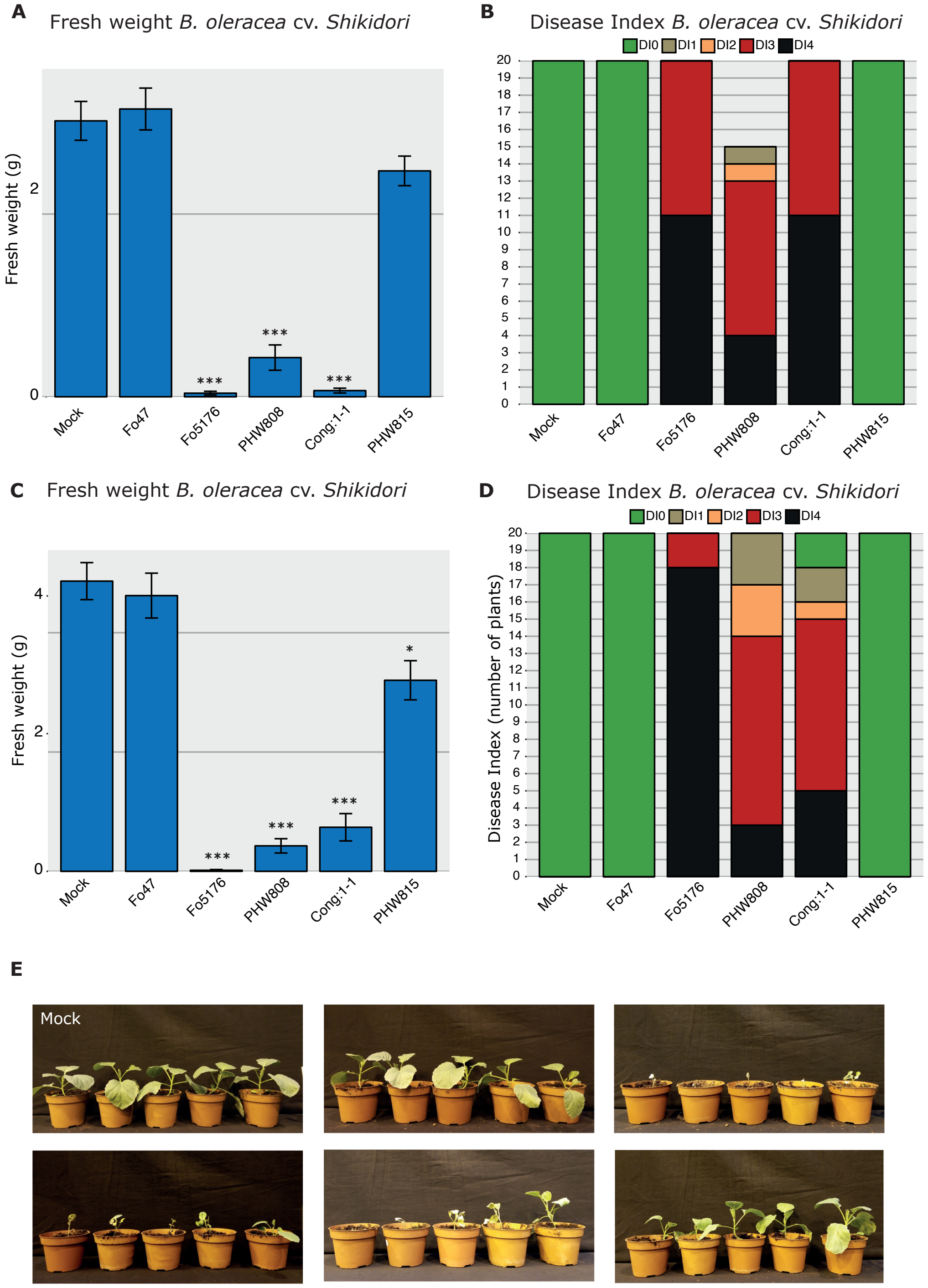


**Supplementary Figure 1: Infection assays of *Fusarium oxysporum* (*Fo*) strains on cabbage (*Brassica oleracea* cv. Shikidori)**. Five representative *Fo* strains were selected including Fo5176. The figure shows two repetitions of the infection assay. Each treatment was applied to 20 plants, except for Fo PHW808 that was applied to 15 plants in the first repetition. A) and C) Fresh weight measurements 13 days after treatment, where bar heights indicate the mean weight and error bars show variation among replicates. Asterisks on top of the bars indicate P-values obtained in a Mann-Whitney test comparing the weights of plants infected with a Fo strain to those of plants that received mock treatment (* < 0.05, ** < 0.01, *** < 0.001). B) and D) Disease index scoring of plants in two independent repetitions. The severity was assessed on a scale from 0 (no disease) to 4 (dead). The height of the bar indicates the number of treated plants. E) Photographs showing representative plants treated with control solution (mock) and different *Fo* strains from the second repetition.

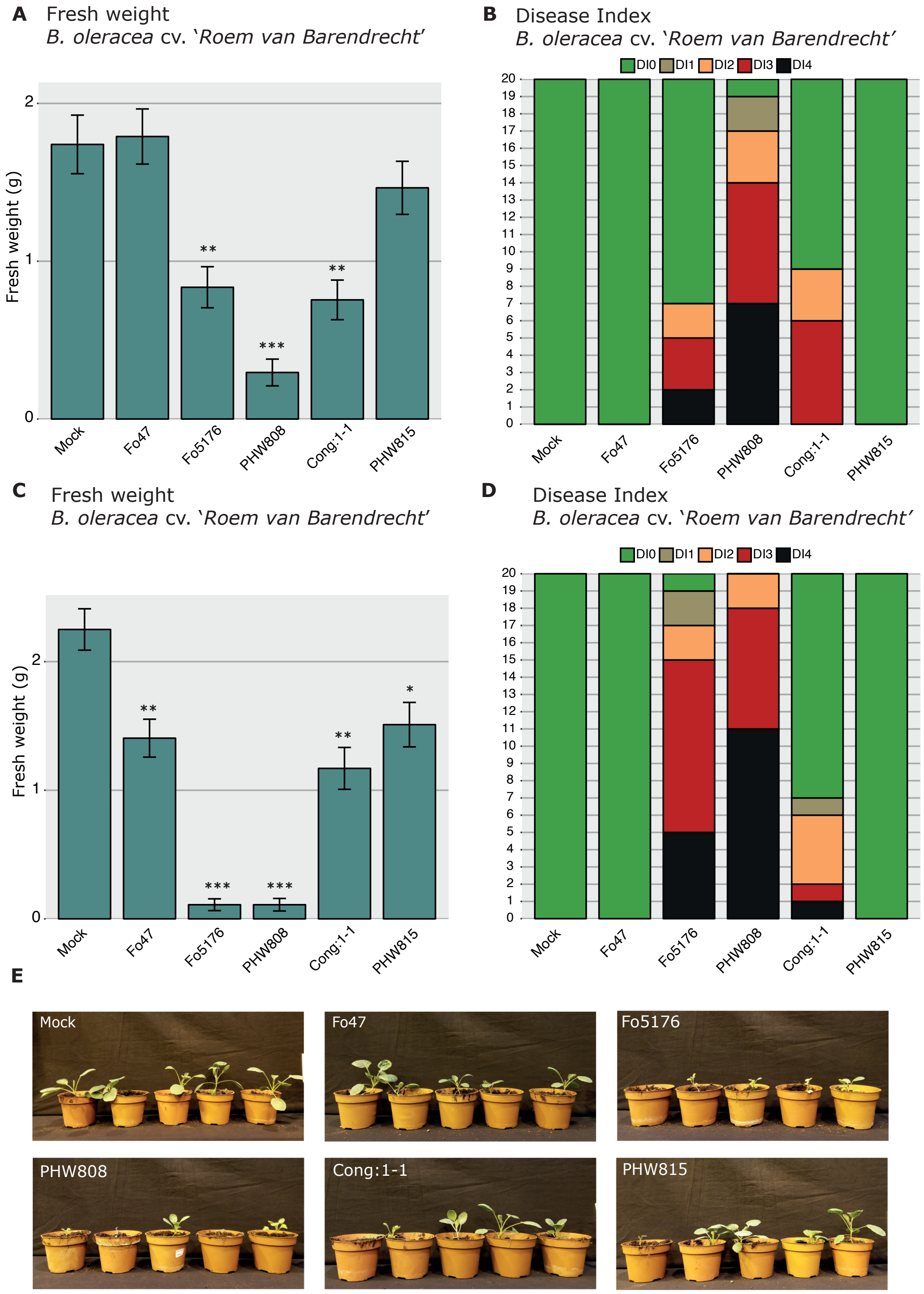


**Supplementary Figure 2: Infection assays of *Fusarium oxysporum* (*Fo*) strains on** Brussels sprouts (*B. oleracea* cv. 'Roem van Barendrecht'). Five representative *Fo* strains were selected including Fo5176. The figure shows two repetitions of the infection assay. Each treatment was applied to 20 plants. A) and C) Fresh weight measurements 13 days after infection, where bar heights indicate the mean weight and error bars show variation among replicates. Asterisks on top of the bars indicate P-values obtained in a Mann-Whitney test comparing the weights of plants infected with a Fo strain to those of plants that received mock treatment (* < 0.05, ** < 0.01, *** < 0.001). B) and D) Disease index scoring of plants in two independent repetitions. The severity was assessed on a scale from 0 (no disease) to 4 (dead). The height of the bar indicates the number of treated plants. E) Photographs showing representative plants treated with control solution () and different *Fo* strains from the second repetition.

### Supplementary Tables

See "Supplementary_Tables.xlsx".

### Supplementary Methods

*RNA extraction and sequencing*

To obtain *in vitro* spores, $10^7$ spores per ml were germinated overnight in ½ Murashige and Skoog Basal Medium (MS; Sigma-Aldrich), 1% sucrose, at 100 rpm. Spores were harvested, washed with sterile water via two centrifugation steps at 4000 g for 15' at 10° C. The pellet was frozen in liquid nitrogen and subsequently freeze-dried before RNA extraction. *In planta* Fo5176 RNA was obtained from hydroponic-grown *A. thaliana* roots two days after fungal inoculation. For, this Col-0 seeds were gas sterilized (50 ml 14% sodium hypochlorite solution and 2 ml 37% HCl) for 4 hours in a sealed bell jar. After 1 hour of gas evaporation, 1 ml of sterile water was added to each tube and the seeds were stratified for 2 days (4° C, dark). The seeds were sown on 2mm foam plugs floating on 330 ml ½ MS + 1% sucrose media at pH 5.7 (adjusted by KOH) in sterile and sealed pots. Plants were grown at 24° C on long-day cycle. The roots were shaded against incoming light and six days after germination the media was exchanged with 330 ml fresh ½ MS. Ten-day-old seedlings were inoculated by adding 20 µl of $10^7$ spores/ml Fo5176 to the media. After 30 min of incubation at 100 rpm on a rotary shaker to prevent spore sinking, the inoculation media was replaced with 330 ml fresh ½ MS media, and the plants grew in the same conditions as before being inoculated. Roots were harvested after 2 days of growth in the same conditions as before inoculation. Roots were harvested by manual removal from foam plugs, dried gently with tissue paper, weighed, and immediately frozen in liquid nitrogen.

For RNA extraction, frozen infected roots and germinated spores were ground with mortar and pestle in liquid nitrogen. A total of 50 -100 µg of ground material was used for RNA extraction using the RNeasy plant mini Kit (Qiagen). Extraction was performed according to the supplier's manual (RNeasy Mini Kit handbook, Fourth edition, June 2012, Qiagen). The extraction buffer RLC with freshly added 10 µl/ml β-mercaptoethanol was used to extract RNA. An on-column DNA digestion (RNase-Free DNase Set, Qiagen) was performed after the first washing step to remove potential DNA contamination from the RNA-sample. Before elution of the RNA, the column was incubated 1 min with 30 µl RNAse-free water to resolve the RNA. RNA concentration was then determined by a Qubit RNA assay (Qubit, Thermo Fisher) and the integrity of the RNA was examined by gel electrophoresis on a 1% agarose gel.

Total mRNA-Seq libraries were prepared using the SENSE mRNA-Seq library Prep Kit V2 (Lexogen) according to the instruction manual with few modifications: 2 µg RNA was used as starting material, the buffer RTL was used for reverse transcription, the incubation time of step 14 was extended to 20 min, the incubation time in step 16 was extended to 2 hours and in step 22, 12 µl of PB was used to purify the synthesized cDNA. The PCR amplification to generate the libraries was conducted with 27 and 24 cycles for the *in planta* sample and the *in vitro* sample, respectively. Purified libraries were stored at -20° C.

All libraries were quality-checked using D1000 ScreenTape on an Agilent 4200 Bioanalyzer at the Functional Genomic Center Zürich (FGCZ). Libraries were pooled based on molarity for sequencing. Sequencing was performed on an Illumina HiSeq2500 sequencer (single-end 125 bp reads) producing ~120 mio reads per sample.

*Soil infection assays on Brassica oleracea*

Fo f. sp. *conglutinans* strains Fo5176 and the biocontrol strain Fo47 were transferred from CDA stock plates to plates with Potato Dextrose Agar and grown at 25° C for four days. Spores of Fo Cong 1-1 Fo PHW808, and Fo f. sp. *raphani* PHW815 that have been stored in 30% glycerol at -80° C, were revived on plates with Potato Dextrose Agar at 25° C for four days. We started a liquid culture by adding a piece of overgrown agar to a flask with 100 ml $NO_3$ medium. Cultures were grown for four days at 25°, 150 RPM. Mycelium was removed by filtering through miracloth. We diluted the resulting spore suspension to obtain a concentration of $5*10^6$ spores per ml. We inoculated 15-20 ten-day old short/long-day grown cabbage (*Brassica oleracea* cv 'Shikidori') and Brussels sprouts (*Brassica oleracea* cv 'Roem van Barendrecht') seedlings by root dipping. Briefly, we carefully removed soil from the seedlings, clipped the root to a length of ca. 1 cm, and placed the seedlings in spore suspension for 2-3 minutes. We performed the same procedure with MilliQ on 20 plants as a negative control. We repotted the seedlings, grow them at 25° C, 65 % humidity, and 16/8 light-dark cycles and scored disease symptoms 13 days after treatment. The disease was scored on a scale of 0 to 4, where 0 = no symptoms, 1 = slight yellowing or swelling of lower leaves, 2 = yellowing of lower leaves, 3 = yellowing and wilting of all leaves, and 4 = plant is dead. The entire experiment was performed twice.
